## Supplementary material for "Role of immigrant males and muzzle contacts in the uptake of a novel food by wild vervet monkeys": All supplementary material referred to in text

### Joint last authorships

#### ***Appendix***

***Appendix 1 – Table 1.*** Table showing details of group membership. Bold text indicates the first to eat in each group. Bold-italic text indicates the first individual to follow the immigrant innovator in extracting and eating peanuts.

| Group | Individual | Age | Sex | Immigration date | Notes | Exposure when first extracting peanuts |
|---|---|---|---|---|---|---|
| $AK_{19}$ | Ati | Adult | M | 03/03/2016 | | n/a |
| $AK_{19}$ | Boc | Adult | M | 15/05/2019 | | n/a |
| $AK_{19}$ | Ghi | Infant | M | n/a | | n/a |

| | | | | | | |
|---|---|---|---|---|---|---|
| AK$_{19}$ | Ghid | Adult | F | n/a | | n/a |
| AK$_{19}$ | Gil | Infant | M | n/a | | n/a |
| AK$_{19}$ | Ginq | Adult | F | n/a | | n/a |
| AK$_{19}$ | Godu | Juvenile | F | n/a | | n/a |
| AK$_{19}$ | Guba | Infant | F | n/a | | n/a |
| AK$_{19}$ | Gubh | Adult | F | n/a | | n/a |
| AK$_{19}$ | Gugu | Adult | F | n/a | | n/a |
| AK$_{19}$ | Guny | Juvenile | F | n/a | | n/a |
| AK$_{19}$ | Guz | Infant | M | n/a | | n/a |
| AK$_{19}$ | Hlu | Juvenile | M | n/a | | n/a |
| AK$_{19}$ | Kek | Adult | M | 31/05/2019 | Unhabituated | n/a |
| AK$_{19}$ | Mat | Juvenile | M | n/a | | n/a |
| AK$_{19}$ | Mbil | Infant | F | n/a | | n/a |
| AK$_{19}$ | Moya | Juvenile | F | n/a | | n/a |
| AK$_{19}$ | Nak | Juvenile | M | n/a | | n/a |
| AK$_{19}$ | Ncok | Infant | F | n/a | | n/a |
| AK$_{19}$ | Nda | Infant | M | n/a | | n/a |
| AK$_{19}$ | Ndaw | Juvenile | F | n/a | | n/a |
| AK$_{19}$ | Ndi | Juvenile | M | n/a | | n/a |
| AK$_{19}$ | Ndik | Juvenile | F | n/a | | n/a |
| AK$_{19}$ | Ndon | Adult | F | n/a | | n/a |
| AK$_{19}$ | Nge | Juvenile | M | n/a | | n/a |
| AK$_{19}$ | Nkos | Adult | F | n/a | | n/a |
| AK$_{19}$ | Nyan | Adult | F | n/a | | n/a |
| AK$_{20}$ | Ghi | Juvenile | M | n/a | | 1 |
| AK$_{20}$ | Ghid | Adult | F | n/a | | 2 |
| AK$_{20}$ | Gil | Juvenile | M | n/a | | 2 |
| AK$_{20}$ | Ginq | Adult | F | n/a | | 3 |
| AK$_{20}$ | Godu | Juvenile | F | n/a | | 2 |
| AK$_{20}$ | Guba | Juvenile | F | n/a | | 3 |
| ***AK$_{20}$*** | ***Gubh*** | ***Adult*** | ***F*** | n/a | | 1 |
| AK$_{20}$ | Gugu | Adult | F | n/a | | 2 |

| | | | | | |
|---|---|---|---|---|---|
| $AK_{20}$ | Guz | Juvenile | M | n/a | 1 |
| $AK_{20}$ | Nak | Juvenile | M | n/a | 2 |
| $AK_{20}$ | Ncok | Juvenile | F | n/a | 1 |
| $AK_{20}$ | Nda | Juvenile | M | n/a | 4 |
| $AK_{20}$ | Ndaw | Juvenile | F | n/a | 3 |
| $AK_{20}$ | Ndik | Juvenile | F | n/a | 2 |
| $AK_{20}$ | Ndon | Adult | F | n/a | 2 |
| $AK_{20}$ | Nge | Juvenile | M | n/a | 2 |
| $AK_{20}$ | Nkos | Adult | F | n/a | n/a |
| $AK_{20}$ | Nyan | Adult | F | n/a | 4 |
| $AK_{20}$ | Twe | Adult | M | 02/04/2020 | 2 |
| **$AK_{20}$** | **Yan** | **Adult** | **M** | **04/03/2020** | 1 |
| BD | Aal | Infant | M | n/a | n/a |
| BD | Aan | Juvenile | M | n/a | 1 |
| BD | Aapi | Adult | F | n/a | n/a |
| BD | Add | Juvenile | M | n/a | 1 |
| BD | Alc | Adult | M | 23/05/2019 | n/a |
| BD | Ard | Infant | M | n/a | n/a |
| BD | Asis | Adult | F | n/a | 3 |
| BD | Bas | Adult | M | 05/05/2017 | n/a |
| BD | Dok | Adult | M | 21/05/2019 | n/a |
| BD | Eina | Adult | F | n/a | n/a |
| BD | Enge | Adult | F | n/a | 4 |
| BD | Fen | Adult | M | 01/09/2017 | 3 |
| BD | Flu | Adult | M | 30/04/2019 | n/a |
| BD | Gese | Adult | F | n/a | n/a |
| BD | Goe | Infant | M | n/a | n/a |
| BD | Han | Adult | M | 06/05/2017 | n/a |
| BD | Hee | Juvenile | M | n/a | 3 |
| BD | Heer | Adult | F | n/a | 4 |
| BD | Hia | Infant | M | n/a | n/a |
| BD | Hibi | Juvenile | F | n/a | n/a |

| | | | | | | |
|---|---|---|---|---|---|---|
| BD | ***Hipp*** | ***Adult*** | ***F*** | n/a | | 1 |
| BD | Hlo | Adult | M | 16/05/2017 | | 1 |
| BD | Hond | Juvenile | F | n/a | | 1 |
| BD | Kom | Adult | M | 2017 | Exact date unknown | n/a |
| BD | Lblind | Adult | F | n/a | | 1 |
| BD | Lewe | Infant | F | n/a | | n/a |
| BD | Miel | Adult | F | n/a | | n/a |
| BD | Mimi | Infant | F | n/a | | n/a |
| BD | Naal | Infant | F | n/a | | n/a |
| BD | Neu | Adult | M | 09/06/2014 | | 1 |
| BD | Non | Infant | M | n/a | | n/a |
| BD | Nooi | Adult | F | n/a | | n/a |
| BD | Numb | Adult | F | n/a | | n/a |
| BD | Nurk | Adult | F | n/a | | n/a |
| BD | Nuu | Juvenile | M | n/a | | n/a |
| BD | Obse | Juvenile | F | n/a | | 1 |
| BD | Oerw | Infant | F | n/a | | n/a |
| BD | Oort | Juvenile | F | n/a | | 1 |
| BD | Ouli | Adult | F | n/a | | 2 |
| BD | Pal | Adult | M | 07/12/2016 | | n/a |
| BD | Pann | Adult | F | n/a | | n/a |
| BD | Piep | Adult | F | n/a | | n/a |
| BD | Pix | Infant | M | n/a | | n/a |
| BD | Poff | Infant | F | n/a | | n/a |
| BD | Pom | Juvenile | M | n/a | | 1 |
| BD | Potj | Adult | F | n/a | | 3 |
| BD | **Pro** | **Adult** | **M** | **13/07/2019** | | 1 |
| BD | Puol | Juvenile | F | n/a | | 1 |
| BD | Rat | Juvenile | M | n/a | | 1 |
| BD | Rede | Adult | F | n/a | | n/a |
| BD | Reen | Infant | F | n/a | | n/a |

| | | | | | |
|---|---|---|---|---|---|
| BD | Rhe | Adult | M | 05/12/2017 | n/a |
| BD | Rid | Infant | M | n/a | n/a |
| BD | Riss | Adult | F | n/a | n/a |
| BD | Sari | Juvenile | F | n/a | n/a |
| BD | Sey | Adult | M | 21/05/2019 | 1 |
| BD | Siel | Adult | F | n/a | n/a |
| BD | Sig | Infant | M | n/a | n/a |
| BD | Sirk | Juvenile | F | n/a | n/a |
| BD | Skem | Infant | F | n/a | n/a |
| BD | Snor | Adult | F | n/a | n/a |
| BD | Spam | Infant | F | n/a | n/a |
| BD | Ted | Adult | M | 25/05/2019 | 1 |
| BD | Ubu | Adult | M | 28/05/2019 | 3 |
| BD | Van | Adult | M | 29/05/2018 | 4 |
| KB | Aar | Infant | M | n/a | 3 |
| KB | Aara | Juvenile | F | n/a | n/a |
| KB | Aare | Adult | F | n/a | n/a |
| KB | Amg | Infant | M | n/a | n/a |
| KB | Amur | Adult | F | n/a | n/a |
| KB | Arn | Juvenile | M | n/a | n/a |
| KB | Lif | Adult | M | 28/04/2015 | n/a |
| KB | Mal | Juvenile | M | n/a | n/a |
| KB | Mara | Adult | F | n/a | n/a |
| KB | Mhao | Juvenile | F | n/a | 4 |
| KB | Mokc | Infant | F | n/a | n/a |
| KB | Nah | Juvenile | M | n/a | n/a |
| KB | Ness | Adult | F | n/a | n/a |
| KB | Yalu | Adult | F | n/a | n/a |
| KB | Yamu | Juvenile | F | n/a | n/a |
| KB | Yara | Infant | F | n/a | n/a |
| KB | Yeni | Adult | F | n/a | n/a |
| KB | Yuko | Juvenile | F | n/a | 4 |

| | | | | | |
|---|---|---|---|---|---|
| KB | Yze | Infant | M | n/a | n/a |
| **LT** | **Bab** | **Adult** | **M** | **24/06/2019** | 1 |
| LT | Ben | Adult | M | 08/06/2018 | 4 |
| LT | Daa | Juvenile | M | n/a | 2 |
| LT | Daen | Adult | F | n/a | 2 |
| LT | Dais | Adult | F | n/a | 2 |
| LT | Dal | Infant | M | n/a | 3 |
| LT | Deli | Adult | F | n/a | 2 |
| LT | Dewe | Juvenile | F | n/a | 2 |
| LT | Dext | Juvenile | F | n/a | 3 |
| LT | Dian | Adult | F | n/a | 1 |
| LT | Digb | Adult | F | n/a | 2 |
| LT | Dil | Juvenile | M | n/a | 4 |
| LT | Dio | Juvenile | M | n/a | 1 |
| LT | Dix | Infant | M | n/a | 1 |
| LT | Dore | Juvenile | F | n/a | 2 |
| LT | Geo | Adult | M | 18/03/2019 | 2 |
| LT | Lail | Infant | F | n/a | n/a |
| LT | Lanc | Adult | F | n/a | 4 |
| LT | Lar | Infant | M | n/a | n/a |
| LT | Laur | Adult | F | n/a | 3 |
| LT | Lava | Juvenile | F | n/a | n/a |
| ***LT*** | ***Lill*** | ***Juvenile*** | ***F*** | n/a | 1 |
| LT | Lizz | Adult | F | n/a | n/a |
| LT | Loui | Juvenile | F | n/a | 4 |
| LT | Rob | Juvenile | M | n/a | n/a |
| **NH** | **Avo** | **Adult** | **M** | **14/05/2018** | 1 |
| NH | Bela | Juvenile | F | n/a | n/a |
| NH | Can | Adult | M | 30/05/2017 | n/a |
| NH | Cus | Adult | M | 21/05/2018 | n/a |
| NH | Gabi | Infant | F | n/a | n/a |
| NH | Gan | Infant | M | n/a | n/a |

| | | | | | |
|---|---|---|---|---|---|
| NH | Gaya | Adult | F | n/a | n/a |
| NH | Gene | Adult | F | n/a | n/a |
| ***NH*** | ***Gran*** | ***Juvenile*** | ***F*** | n/a | 1 |
| NH | Guat | Juvenile | F | n/a | n/a |
| NH | Jixi | Juvenile | M | n/a | n/a |
| NH | Lima | Juvenile | F | n/a | 4 |
| NH | Prai | Juvenile | F | n/a | n/a |
| NH | Pret | Adult | F | n/a | n/a |
| NH | Pro | Juvenile | M | n/a | n/a |
| NH | Pru | Juvenile | M | n/a | n/a |
| NH | Pye | Infant | M | n/a | n/a |
| NH | Raba | Juvenile | F | n/a | n/a |
| NH | Renn | Adult | F | n/a | n/a |
| NH | Rev | Infant | M | n/a | n/a |
| NH | Reva | Adult | F | n/a | n/a |
| NH | Rey | Juvenile | M | n/a | n/a |
| NH | Rioj | Infant | F | n/a | n/a |
| NH | Roma | Adult | F | n/a | n/a |
| NH | Rosl | Juvenile | F | n/a | n/a |
| NH | Twe | Adult | M | 24/05/2014 | n/a |
| NH | Ula | Juvenile | M | n/a | n/a |
| NH | Umt | Juvenile | M | n/a | n/a |
| NH | Upps | Adult | F | n/a | 4 |
| NH | Ura | Infant | M | n/a | n/a |
| NH | Wol | Adult | M | 29/05/2018 | n/a |
| NH | Xal | Infant | M | n/a | 4 |
| NH | Xala | Adult | F | n/a | n/a |
| NH | Xia | Juvenile | M | n/a | 1 |
| NH | Xin | Infant | M | n/a | n/a |
| NH | Xian | Adult | F | n/a | n/a |
| NH | Yan | Adult | M | 14/05/2019 | n/a |

19

***Appendix 1 – Table 2. Dominance hierarchies for all groups were significantly linear.***

| Group | AK19 | AK20 | BD | KB | LT | NH |
|---|---|---|---|---|---|---|
| h’ | 0.45 | 0.26 | 0.21 | 0.53 | 0.22 | 0.33 |
| p-value | <0.001 | <0.001 | <0.001 | <0.001 | 0.017 | <0.001 |
